# Integrated systems biology identifies disruptions in mitochondrial function and metabolism as key contributors to heart failure with preserved ejection fraction (HFpEF)

**DOI:** 10.1101/2024.10.25.619450

**Authors:** Andrew A. Gibb, Kyle LaPenna, Ryan B. Gaspar, Nadina R. Latchman, Yinfei Tan, Carmen Choya-Foces, Jake E. Doiron, Zhen Li, Huijing Xia, Michael P. Lazaropoulos, Mariell Conwell, Thomas E. Sharp, Traci T. Goodchild, David J. Lefer, John W. Elrod

## Abstract

**Background:** Heart failure with preserved ejection fraction (HFpEF) accounts for ∼50% of HF cases, with no effective treatments. The ZSF1-obese rat model recapitulates numerous clinical features of HFpEF including hypertension, obesity, metabolic syndrome, exercise intolerance, and LV diastolic dysfunction. Here, we utilized a systems-biology approach to define the early metabolic and transcriptional signatures to gain mechanistic insight into the pathways contributing to HFpEF development.

**Methods:** Male ZSF1-obese, ZSF1-lean hypertensive controls, and WKY (wild-type) controls were compared at 14w of age for extensive physiological phenotyping and LV tissue harvesting for unbiased metabolomics, RNA-sequencing, and assessment of mitochondrial morphology and function. Utilizing ZSF1-lean and WKY controls enabled a distinction between hypertension-driven molecular changes contributing to HFpEF pathology, versus hypertension + metabolic syndrome.

**Results:** ZSF1-obese rats displayed numerous clinical features of HFpEF. Comparison of ZSF1-lean vs WKY (i.e., hypertension-exclusive effects) revealed metabolic remodeling suggestive of increased aerobic glycolysis, decreased β-oxidation, and dysregulated purine and pyrimidine metabolism with few transcriptional changes. ZSF1-obese rats displayed worsened metabolic remodeling and robust transcriptional remodeling highlighted by the upregulation of inflammatory genes and downregulation of the mitochondrial structure/function and cellular metabolic processes. Integrated network analysis of metabolomic and RNAseq datasets revealed downregulation of nearly all catabolic pathways contributing to energy production, manifesting in a marked decrease in the energetic state (i.e., reduced ATP/ADP, PCr/ATP). Cardiomyocyte ultrastructure analysis revealed decreased mitochondrial area, size, and cristae density, as well as increased lipid droplet content in HFpEF hearts. Mitochondrial function was also impaired as demonstrated by decreased substrate-mediated respiration and dysregulated calcium handling.

**Conclusions:** Collectively, the integrated omics approach applied here provides a framework to uncover novel genes, metabolites, and pathways underlying HFpEF, with an emphasis on mitochondrial energy metabolism as a potential target for intervention.

## INTRODUCTION

Heart failure (HF) is a growing epidemic. In the U.S. alone, >6.7 million people over the age of 20 have HF and this is projected to increase to >8.5 million people by 2030^1,2^. Nearly one-quarter of people will develop HF in their lifetime^1–3^ and current HF mortality rates are higher today than in 1999^4^. Of those diagnosed with HF, ∼50% have heart failure with preserved ejection fraction (HFpEF)^1–3,5^. HFpEF patients present with elevated left ventricular (LV) filling pressure despite normal LV ejection fraction (≥50%). At present there are very limited treatments for HFpEF, and the 5-year mortality rate is merely 50%^1–3^. Clinical trials of drugs that are effective in HF with reduced ejection fraction (HFrEF) have uniformly failed in HFpEF^1,3^. Due to our limited understanding of mechanisms which drive HFpEF and lack of therapeutic strategies to treat this devastating disease, the NIH-NHLBI has issued a statement of emphasis detailing the research priority of HFpEF and identified HFpEF as the greatest unmet need in cardiovascular medicine^6^.

While clinicians struggle to treat HFpEF patients, research scientists have grappled with preclinical models to study the pathobiology of HFpEF^7–12^ to improve our understanding of this complex, multi-organ disease. Clinically relevant models are required to fully elucidate molecular disease mechanisms and effectively translate new therapies from bench to bedside. Towards this end, the (ZSF1) rat has been proposed as an animal model for HFpEF^13,14^. This model was created by crossing rat strains with two separate leptin receptor mutations (fa and facp), the lean female Zucker diabetic fatty (ZDF) rat (+/fa) and the lean male spontaneously hypertensive heart failure (SHHF) rat. Offspring homozygous for both mutations (fa:facp) create a hybrid rat with central obesity and hypertension (ZSF1-Obese rat) resulting in spontaneous cardiometabolic HFpEF whereas the heterozygous lean offspring (ZSF1-Lean rat) exhibit no signs of obesity and diabetes^15^. Previous studies have shown that obese ZSF1 rats develop significant diastolic dysfunction between 10-20 weeks of age with concentric LV remodeling and hypertrophy like that observed in HFpEF patients^16^. In addition to LV diastolic dysfunction, previous studies have demonstrated skeletal muscle pathology, exercise intolerance, endothelial dysfunction, systemic inflammation, and renal and hepatic abnormalities that are consistent with cardiometabolic HFpEF^13,16–19^. We have previously demonstrated that the ZSF1 rat is responsive to therapeutic interventions when delivered early during the progression of HFpEF^18,19^. The severity of HFpEF in terms of cardiometabolic pathology has been shown to be similar between male and female ZSF1 obese rats^20^, which is not the case for the popular “two-hit” mouse model of HFpEF in which female mice are protected against the development of HFpEF^21^. In summary, the ZSF1 obese rat represents a clinically-relevant and superior model for the elucidation of novel mechanisms responsible for the development and progression of HFpEF.

To uncover potentially novel and critical mechanisms in HFpEF, we provide an in-depth characterization the ZSF1 obese rat model of HFpEF using several physiological, biochemical, molecular, and omics approaches. We evaluated male ZSF1-obese, ZSF1-lean hypertensive controls, and WKY (wild-type) lean, normotensive controls at an early stage in the development of HFpEF (14-wks of age), performing extensive physiological phenotyping in conjunction with unbiased metabolomics and transcriptomics. Our results reveal that the addition of obesity/metabolic syndrome upon hypertension and vascular dysfunction is a primary contributor to gross cardiac transcriptional and metabolic remodeling, driving the development of HFpEF. Most notably, mitochondrial energy metabolism pathways were highly disrupted resulting in an energetic deficit that correlated with maladaptive mitochondrial ultrastructural remodeling and functional impairment. These findings support an integrated framework to identify metabolic and transcriptional pathways that are disrupted in, and contribute to, HFpEF progression that will optimally yield new therapeutic targets.

## METHODS

### Experimental Animals

Wistar Kyoto (WKY), ZSF1-lean, and ZSF1-obese male rats were purchased from Charles River laboratories and used in all experiments contained within this study (n=5 to 7 per group). Animals were purchased and held at Temple University Lewis Katz School of Medicine (TU-LKSOM) or LSU Health Sciences Center (LSUHSC) in a temperature controlled and 12-hour light/dark cycle for the entirety of studies. All studies were approved by TU-LKSOM and LSUHSC Institutional Animal Care and Use Committees (IACUC) and received animal care at TU-LKSOM and LSUHSC according to the Association for Assessment and Accreditation of Laboratory Animal International (AAALAC) guidelines.

### Study Design

Both ZSF1-lean, ZSF1-obese, and WKY controls were investigated at 14 wks of age. Physiologic parameters of body weight, transthoracic echocardiography, and exercise capacity testing are as described below. Further investigation into pathophysiology of these separate animal models was performed utilizing left ventricular (LV) and systemic invasive hemodynamic measurements along with ex vivo assessments of mitochondrial ultrastructure and function. Isolated cardiac LV tissue samples were also submitted for RNAseq and unbiased metabolomics.

### Echocardiography

Transthoracic echocardiography of all groups was performed with a Vevo 2100 echocardiography system (FUJIFILM VisualSonics). Left ventricular diastolic measurements were performed using an apical four chambered view of the heart. Left ventricular systolic measurements were performed using a long-axis view. Animals were anesthetized using inhaled isoflurane at an induction dose of 3% with a maintenance dose of 1% for the longevity of the experiment. Heart rate was maintained at approximately 250-300 beats per minute (BPM) for the data collection period as previously described^22^.

### Exercise Capacity Testing

ZSF1-lean, ZSF1-obese, and WKY control rats were assessed for exercise intolerance utilizing a IITC Life Science 800 Series treadmill. Animals were first acclimated to the treadmill for a period of 5 minutes with no movement, they were then brought through a warmup phase consisting of initially 6 meters per minute which was thereby increased to 12 meters per minute for a 4-minute ramp up time, for a total warmup phase of 5 minutes. For data collection as presented, the animals were run at a rate of 12 meters per minute with 0° incline until exhaustion, which was defined as animal placement on the shock pads for more than 3 seconds. Exercise capacity was then determined by the total distance run.

### Terminal Invasive Hemodynamics and Sacrifice

At 14 wks of age, animals were anesthetized via inhaled isoflurane at a concentration of 3% for induction and 1% maintenance during the following procedure. The rodent neck and associated structures were dissected for exposure of the common carotid which was canulated with a 1.2 F high-fidelity pressure catheter, measuring the systemic pressures at systole and diastole accordingly for multiple cardiac cycles. The pressure catheter was then carefully introduced into the left ventricle of the animal. Left ventricular end diastolic pressures (LVEDP) and ventricular relaxation time constant (Tau) were measured after multiple cardiac cycles to obtain an average measurement. The catheter was then removed, and the rat was subsequently exsanguinated and sacrificed with tissues and plasma harvested for additional measurements as previously described^23^.

### Mitochondrial function

Heart mitochondria were isolated and subjected to respiratory function assays using the Seahorse XF96, like that described previously^7,24^. Briefly, ∼100 mg left ventricular heart pieces were washed 5× with cold buffer A (220 mM mannitol, 70 mM sucrose, 5 mM MOPS, 1 mM EDTA; pH 7.2 with KOH) followed by homogenization using a glass-col homogenizer in 2 ml of buffer A containing 0.2% fatty acid-free BSA. Homogenate was then subjected to centrifugation at 800 × *g* for 10 min followed by supernatant collection and centrifugation at 10,000 × *g* for 10 min. The pellet containing mitochondria was then resuspended in 1 ml fresh buffer A (without BSA) and centrifuged at 10,000 × *g*, with this step repeated once. The washed mitochondrial pellet was then resuspended in 150 µl respiration buffer (120 mM KCl, 25 mM Sucrose, 10 mM HEPES, 1 mM MgCl_2_, 5 mM KH_2_PO_4_; pH 7.2 with KOH) and kept on ice.

To determine mitochondrial function, samples were diluted to a concentration of 2.5 µg (protein) in 50 µl respiration buffer per well and centrifuged onto XF96 microplates at 500 × *g* for 3 min at 4°C. State 3 respiration in response to substrates were measured after injection of pyruvate + malate (5.0 mM + 2.5 mM, final concentrations) or succinate + rotenone (10 mM + 1 µM, final concentrations) to assess complex I and II rates, respectively. Fatty acid oxidation was assessed in response to palmitoyl-l-carnitine + malate (50 µM + 2.5 mM, final concentration). The oxygen consumption rates recorded after injection of oligomycin (1 µg/ml), an inhibitor of ATP synthase, served as a measure of State 4 respiration. Following State 4 respiratory measurements, injection of FCCP, a mitochondrial uncoupler, provided ETC complex maximal respiratory capacity. Respiratory control ratios, state 3/state 4, were calculated as a measure of the coupling of oxygen consumption to ATP production.

### Mitochondrial calcium uptake assay

Isolated mitochondria were diluted in Isolated <u>M</u>itochondria <u>A</u>ssay <u>B</u>uffer (IMAB; 125 mM KCl, 10 mM NaCl, 20 mM HEPES, 2 mM MgCl_2_, 2 mM KH_2_PO_4_, pH 7.2 with KOH). Mitochondria were loaded into 96-well plates (final concentration of 1 µg/µL), supplemented with 10 mM succinate (Sigma-Aldrich, S3674), 10 mM malate (Sigma-Aldrich, 240176) and 10 mM pyruvate (Sigma-Aldrich, P5280), and 1 µM calcium green-5N hexapotassium salt (Invitrogen, C-3737). Final volume at the start of the assay was 50 µL. Fluorescence was measured every 200 ms at 506 nm_ex_/532 nm_em_ using a TECAN Infinite M1000 Pro plate reader set at 37°C. After 120 sec of baseline measurements, successive injections of 2.5 µM CaCl_2_ (5 µL of 25 µM CaCl_2_ stock prepared in IMAB) were administered every 120 sec. To generate a standard curve of extramitochondrial Ca_2+_ (bath concentration), the same experimental setup was employed without addition of mitochondria to the well. The standard curve was utilized to calculate the extramitochondrial calcium remaining post mitochondrial uptake (average of last 100 sec per injection cycle) and to determine the percent mitochondrial calcium uptake following successive injections. All methods are as described^25–27^.

### Mitochondrial swelling assays

Isolated mitochondria were diluted in Isolated <u>M</u>itochondria <u>A</u>ssay <u>B</u>uffer (IMAB; 125 mM KCl, 10 mM NaCl, 20 mM HEPES, 2 mM MgCl_2_, 2 mM KH_2_PO_4_, pH 7.2 with KOH). Mitochondria were loaded into 96-well plates (final concentration of 1 µg/µL), supplemented with 10 mM succinate (Sigma-Aldrich, S3674), 10 mM malate (Sigma-Aldrich, 240176) and 10 mM pyruvate (Sigma-Aldrich, P5280). Final volume was 150 µL per well. Absorbance was measured every 5 seconds at 540 nm using a TECAN Infinite M1000 Pro plate reader set at 37°C with plate shaking between measurements. After 2 minutes of baseline measurements, a single Ca^2+^ bolus of 500 µM CaCl_2_ (7.5 µL of 10 mM CaCl_2_ stock prepared in IMAB) was administered with measurements recorded every 5 sec for 10 min. All methods are as described^25–27^.

### Transmission Electron Microscopy (TEM)

Left ventricle tissue cut to ∼3 mm^3^ were fixed in 2% PFA + 2% glutaraldehyde in 0.1 M sodium cacodylate buffer, pH 7.4, and stored at 4°C for 48 h. Tissues were washed 3x for 15 min each in 0.1 M sodium cacodylate buffer, pH 7.4, and then post-fixed in freshly prepared 1.5% potassium ferrocyanide and 1% osmium tetroxide in 0.1 M sodium cacodylate buffer pH 7.4 for 2 h. The samples were washed with water 4x for 15 min each followed by *en bloc* staining overnight with 1% uranyl acetate (aq). Following washing 3x with H_2_O for 15 min each, tissues were dehydrated in an ascending acetone series (25% acetone, 50% acetone, 75% acetone, 95% acetone, 100% anhydrous acetone, 100% anhydrous acetone), 15 min each step. Samples were infiltrated with Spurr’s resin (25% resin in acetone, 50% resin in acetone, 75% resin in acetone, 100% resin, 100% resin), 1 h each step followed by overnight incubation in 100% Spurr’s resin. The next day, one last exchange in 100% Spurr’s resin was performed before samples were placed in aluminum weigh dishes with fresh resin and polymerized at 60°C overnight. Following polymerization, tissues in proper orientation were excised from the resin with a jeweler’s saw and glued onto supports. Muscle tissues were sectioned with a Leica UC7 ultramicrotome, and 60 nm thick sections were collected onto 200 mesh copper grids with a formvar-carbon support film. Grids were post-stained with 2% uranyl acetate in 50% methanol and Reynolds lead citrate. Grids were examined and imaged in a FEI Tecnai 12 120 keV digital TEM, with images acquired at various magnifications (e.g. 1,100x – 21,000x).

### Morphometric analysis of TEM images

Analysis of mitochondria, lipid droplets (LD), LD-mitochondria associations, and sarcomere lengths were performed using ImageJ/FIJI (NIH). After calibration for distance, shape descriptors and size measurements were obtained by manually tracing only discernable mitochondria or lipid droplets. Circularity is computed as [4pi × (area/perimeter^2^)] and roundness is computed as [4 pi × (surface area)/(pi × major axis^2^); values of 1 indicate perfect spheres. Feret Diameter represents the longest distance between any two points within a given mitochondrion^28^. A custom Phyton plugin (MitoCareTools) was adopted for quantification of lipid-mitochondrion associations^29,30^. Areas where the LD was <100nm from the outer mitochondrial membrane (OMM) were determined as a LD-mitochondrion interface. To obtain mean gap distance, the LD membrane was first traced followed by tracing the mitochondrion OMM with values obtained from the plugin.

### Protein Immunoblotting

Remaining isolated mitochondria from our calcium and respiratory assays were pelleted and lysed in RIPA buffer supplemented with phosphatase inhibitors (Roche, 4906837001) and protease inhibitors (Sigma, S8830). Samples were kept on ice for 30 min with agitation via vortex every 10 min. Samples were then centrifuged at 13,000 × *g* for 20 min at 4°C. The supernatant was collected, and protein concentration quantified using the Pierce 660nm Protein Assay Reagent (Thermo Fisher Scientific). Equal amounts of protein (5 ug) were run by gel electrophoresis on polyacrylamide Tris-glycine SDS gels. Gels were transferred to PVDF (EMD Milipore, IPFL00010) and membranes were blocked for 1 h in Blocking Buffer (Rockland, MB-070) followed by incubation with primary antibody overnight at 4°C on a rocker. Membranes were then washed in TBS-T 3x for 5 min each and incubated in a fluorescent secondary antibody for 1 h at RT. Membranes were then washed in TBS-T 3x for 5 min each and imaged on a Licor Odyssey system. Antibodies in the study were used at a concentration of 1:1000 and include: VDAC1/3 (Abcam, ab14734), MCU (Cell Signal, 14997), MICU1 (Novus Bio, BP1-86663), MICU2 (Novus Bio, BP2-92063), MCUB (Sigma Aldrich, HPA024771), Total OxPHOS Cocktail (Abcam, ab110413).

### RNA sequencing

Left ventricular heart pieces were immediately flash frozen in liquid N_2_ following excision and subjected to RNAseq analysis. Total RNA was isolated using a fibrous tissue RNA isolation kit (Qiagen). The TrueSeq stranded mRNA library prep kit was used to enrich polyA mRNAs via poly-T based RNA purification beads which were then amplified using HiSeq rapid SR cluster kit and multiplexed and run using the HiSeq rapid SBS kit. Reading depth was ∼30M reads per sample and single-end 75 bp fragments were generated for bioinformatic analysis. All kits for sequencing were obtained from Illumina and all sequencing was performed on the Illumina HiSeq2500 sequencer. RNA transcripts were aligned to the *Rnor_6.0* assembly using HISAT2 v2.1.0 and quantified using HTSeq v0.11.2. Differential expression analysis was performed between groups using DESeq2 v1.22.2. Genes were considered differentially expressed when they met a fold change ≥2.0 and FDR ≤0.05. Gene ontology (GO) analysis was accomplished using DAVID GO analysis tools. All RNA-sequencing data will be submitted to the GEO repository with the appropriate accession # at time of publication.

### Metabolomic analysis

Left ventricular heart pieces were immediately flash frozen in liquid N_2_ following excision to most accurately capture the *in vivo* cardiac metabolome. Samples were prepared by Metabolon using their automated MicroLab STAR^®^ system (Hamilton Company, Reno, NV). First, tissue homogenates were made in water at a ratio of 5 µL per mg of tissue. For quality control, several recovery standards were added prior to the first step in the extraction process. To remove protein, dissociate small molecules bound to protein or trapped in the precipitated protein matrix, and to recover chemically diverse metabolites, proteins were then precipitated with methanol (final concentration 80% v/v) under vigorous shaking for 2 min (Glen Mills GenoGrinder 2000) followed by centrifugation. For quality assurance and control, a pooled matrix sample was generated by taking a small volume of each experimental sample to serve as a technical replicate throughout the data set. Extracted water samples served as process blanks. A cocktail of standards known not to interfere with the measurement of endogenous compounds was spiked into every analyzed sample, allowing instrument performance monitoring and aiding chromatographic alignment.

The extract was divided into fractions for analysis by reverse phase (RP)/UPLC-MS/MS with positive ion mode electrospray ionization (ESI), by RP/UPLC-MS/MS with negative ion mode ESI, and by HILIC/UPLC-MS/MS with negative ion mode ESI. Samples were placed briefly on a TurboVap^®^ (Zymark) to remove the organic solvent. All methods utilized a Waters ACQUITY UPLC and a Thermo Scientific Q-Exactive high resolution/accurate mass spectrometer interfaced with a heated electrospray ionization (HESI-II) source and Orbitrap mass analyzer operated at 35,000 mass resolution. The sample extract was reconstituted in solvents compatible with each MS/MS method.

Each reconstitution solvent contained a series of standards at fixed concentrations to ensure injection and chromatographic consistency. One aliquot was analyzed using acidic positive ion conditions, chromatographically optimized for hydrophilic compounds. In this method, the extract was gradient eluted from a C18 column (Waters UPLC BEH C18-2.1×100 mm, 1.7 µm) using water and methanol, containing 0.05% perfluoropentanoic acid (PFPA) and 0.1% formic acid (FA). For more hydrophobic compounds, the extract was gradient eluted from the C18 column using methanol, acetonitrile, water, 0.05% PFPA and 0.01% FA. Aliquots analyzed using basic negative ion optimized conditions were gradient eluted from a separate column using methanol and water, containing 6.5 mM ammonium bicarbonate (pH 8). The last aliquot was analyzed via negative ionization following elution from a HILIC column (Waters UPLC BEH Amide 2.1×150 mm, 1.7 µm) using a gradient consisting of water and acetonitrile with 10 mM ammonium formate (pH 10.8). The MS analysis alternated between MS and data-dependent MS^n^ scans using dynamic exclusion. The scan range covered 70–1000 *m/z*.

Raw data were extracted, peak-identified and processed using Metabolon’s proprietary hardware and software. Compounds were identified by comparison to library entries of purified, authenticated standards or recurrent unknown entities, with known retention times/indices (RI), mass to charge ratios (*m/z*), and chromatographic signatures (including MS/MS spectral data). Biochemical identifications were based on three criteria: retention index within a narrow RI window of the proposed identification, accurate mass match to the library±10 ppm, and the MS/MS forward and reverse scores between experimental data and authentic standards. Proprietary visualization and interpretation software (Metabolon, Inc., Durham, NC) was used to confirm the consistency of peak identification among the various samples. Library matches for each compound were checked for each sample and corrected, if necessary. Area under the curve was used for peak quantification.

Original scale data (raw area counts) were analyzed using Metaboanalyst 5.0 software (http://www.metaboanalyst.ca/). Metabolites with greater than 50% of the values missing were omitted from the analysis, and missing values were imputed by introducing values with 1/5 of the minimum positive value of each variable. An interquartile range filter was used to identify and remove variables unlikely to be of use when modeling the data. The data were log-transformed and auto-scaled (mean-centered and divided by the standard deviation of each variable). Univariate (e.g., volcano plots) and multivariate (e.g., PCA) analyses were then performed. For multiple comparison testing, *q* (FDR) values were calculated in R using a method embedded within the Metaboanalyst software, controlling for the false discovery rate. Metabolites were considered significantly different when they met a fold change ≥1.25 and FDR ≤0.05.

### Integrated Pathway Network Analyses

Integrated network analyses utilizing both the transcriptomic and metabolomic datasets were performed using Metaboanalyst 5.0 software. Integrated pathway maps were generated using BioRender.

### Statistical Analysis

Statistical analysis was performed using GraphPad Prism 9, Metaboanalyst, and the R program. Statistical parameters including the value of n (number of cats), the definition of center, dispersion and precision measures (mean±SEM or SD), and statistical significance is reported in the figures and figure legends. A *P* value of ≤0.05 was considered statistically significant. For the metabolomics and transcriptomics data sets, an FDR value of ≤0.05 was considered statistically significant. For direct comparisons, statistical significance was calculated by unpaired or paired Student *t* test. Details on the statistical methods employed for the metabolomics and RNA-seq data sets can be found within their respective methods sections.

## RESULTS

### The clinical features of HFpEF are recapitulated in the ZSF1-Obese rat

We investigated whether the ZSF1-Obese rat, which is both hypertensive and obese, phenocopies the clinical characteristics of HFpEF and aimed to identify potential molecular and metabolic mechanisms contributing to HFpEF (**Fig. 1A**); ZSF1-Lean (hypertensive lacking obesity/metabolic syndrome) and WKY rats were included as controls. ZSF1-Obese rats demonstrated a 60% and 40% increase in body weight compared to WKY and ZSF1-Lean controls, respectively (**Fig. 1B**). Both Lean and Obese rats were hypertensive, with elevated systolic (∼155 mmHg) and diastolic (∼110 mmHg) blood pressures (**Fig. 1C**). Distance run on a treadmill was 83% less in ZSF1-Obese rats, indicating severe exercise intolerance, which was also observed in lean rats (**Fig. 1D**). Echocardiography revealed a significant elevation in the E/e’ in ZSF1-Obese rats with preserved ejection fraction (EF%) **(Fig. 1E,F**). Invasive hemodynamics (PV Loop) indicated a 6-fold increase in LVEDP (left ventricular end diastolic pressure; **Fig. 1E**), a hallmark feature distinguishing HFpEF from HFrEF^31^. Left ventricular, atrial, liver, and kidney weights when normalized to tibia length were greatest in ZSF1-Obese rats vs. controls (**Fig. 1G, Supplemental Fig. 1**), indicating tissue hypertrophy and/or edema. Collectively, the ZSF1-Obese rat displays numerous features of clinical HFpEF including obesity, hypertension, exercise intolerance, diastolic dysfunction with preserved ejection fraction, and cardiac hypertrophy.

**Fig. 1:**
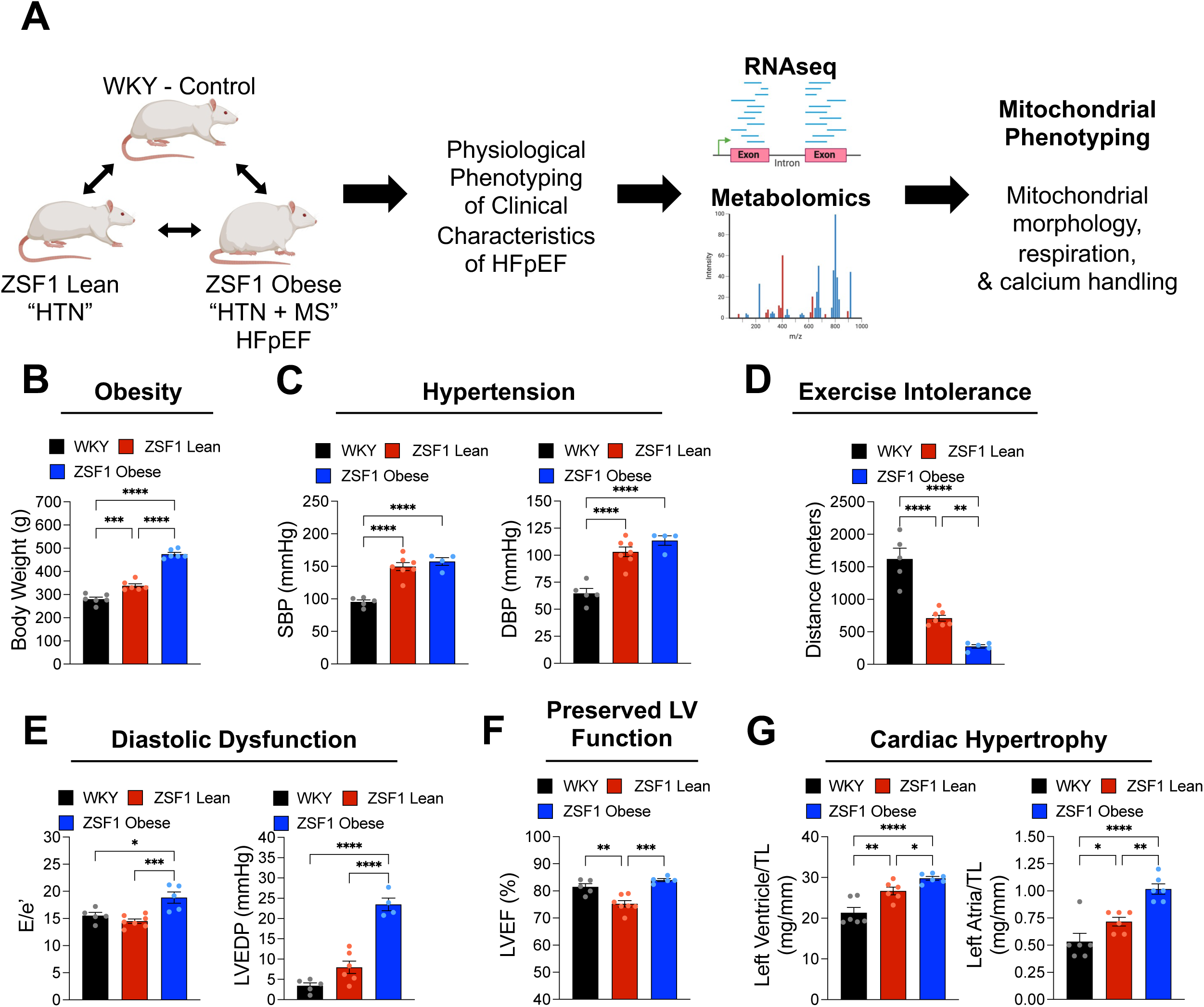
Clinical manifestations of HFpEF are observed in the ZSF1-Obese rat. Physiological characterization of HFpEF. (**A**) Schematic of study design. (**B**) Body weights of WKY (control), Lean (Hypertensive; HTN), and ZSF1-Obese (Metabolic Syndrome + HTN; HFpEF) rats. (**C**) Assessment of systolic (SBP) and diastolic (DBP) blood pressure obtained during invasive hemodynamics. (**D**) Distance run during a treadmill exercise capacity test. (**E**) Indices of cardiac left ventricular diastolic function assessed by echocardiography for the E/e’ ratio and invasive hemodynamics for the left ventricular end diastolic pressure (LVEDP). (**F**) Determination of cardiac systolic function assessed by the echocardiography for the left ventricular ejection fraction (LVEF%). (**G**) Gravimetric assessment of left ventricle and left atria normalized to tibia length (TL). n = 4-7 male rats per group, mean ± SEM. One-way ANOVA with Holm-Sidak’s post-hoc test, *p ≤ 0.05, **p ≤ 0.01, ***p ≤ 0.001, ****p ≤ 0.0001.

### Lean hypertensive rats demonstrate significant metabolic remodeling with few transcriptional changes

To identify potential molecular and metabolic pathways contributing to disease development, RNAseq and quantification of the steady-state abundance of metabolites was performed in hearts from all 3 genotypes, initially assessing those changes mediated by hypertension alone by comparing ZSF1-Lean to WKY controls. We observed differential expression of 233 genes in Lean hearts, with 149 increased and 84 decreased in expression (fold change [FC] ≥ 2.0 and FDR ≤ 0.05) (**Fig. 2A**). Gene ontology (GO) analysis of the differentially expressed transcripts surprisingly revealed no significant enrichment of biological or KEGG pathways (**Fig. 2B-E**), suggesting diffuse and non-specific transcriptional remodeling. Metabolomics analysis identified 120 metabolites increased in abundance and 85 decreased in abundance (FC ≥ 1.25 and FDR ≤ 0.05) (**Fig. 2A**). Unlike our transcriptomics dataset, pathway enrichment analysis of the cardiac metabolome revealed significant changes (*p* < 0.05) in nucleotide metabolism, amino acid metabolism, and pathways critical for energy metabolism (e.g., glycolysis, pyruvate, Krebs cycle) (**Fig. 2F**). Collectively, these results suggest that chronic hypertension alone is sufficient to robustly remodel cardiac metabolism while minimally impacting the transcriptome.

**Fig. 2:**
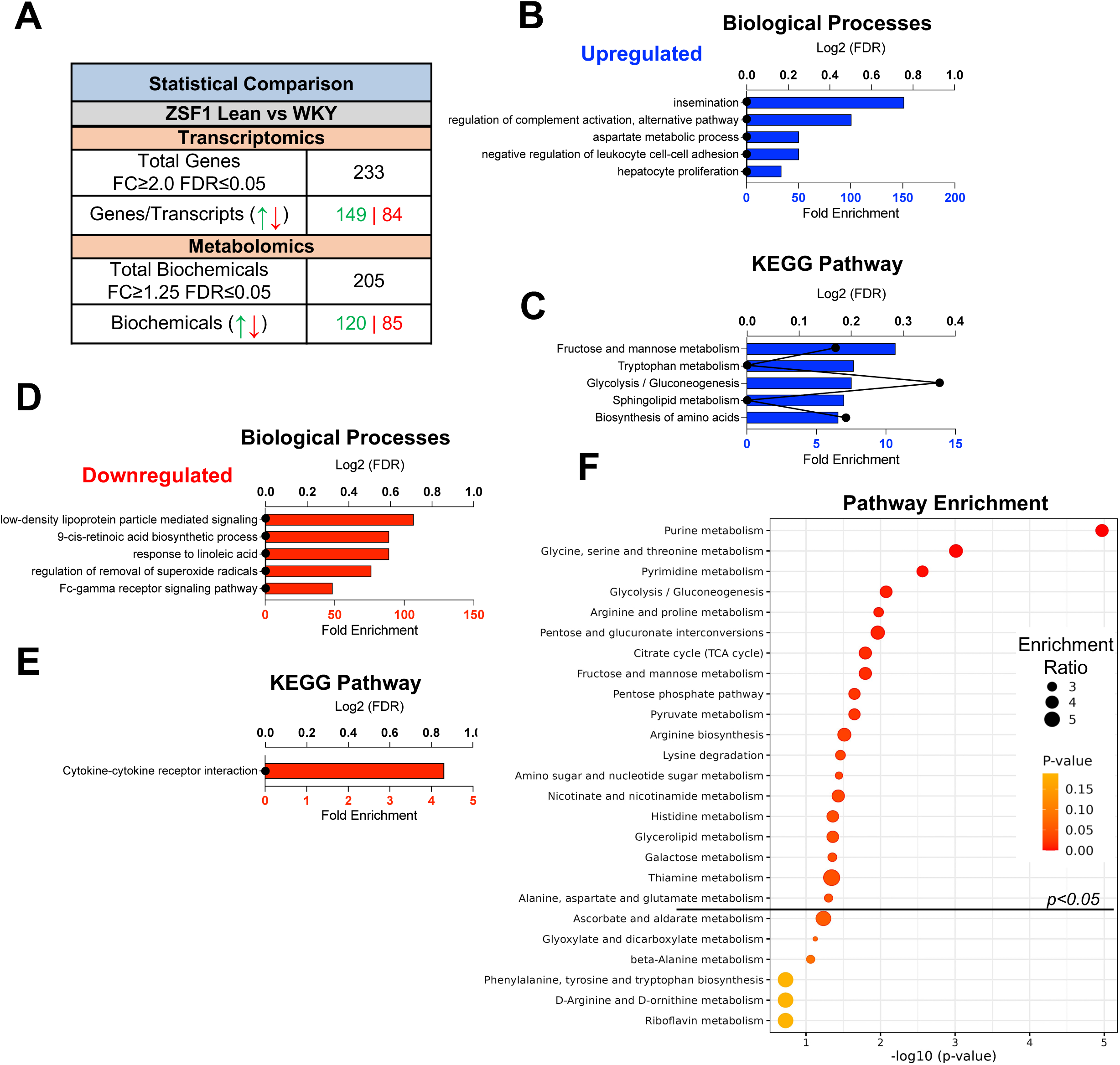
Hypertension significantly impacts the cardiac metabolome with minimal impact on the transcriptome. RNAseq and metabolomic comparisons of hearts from Lean vs WKY control rats. (**A**) Summary of the upregulated and downregulated transcriptional and metabolic changes. Gene ontology analysis revealing the top upregulated (**B**) biological processes and (**C**) KEGG pathways of those genes found to be differentially expressed. Gene ontology analysis revealing the top downregulated (**D**) biological processes and (**E**) KEGG pathways of those genes found to be differentially expressed. (**F**) Pathway enrichment analysis of the cardiac metabolome indicated those pathways found to be most significantly affected in ZSF1 rats due to hypertension. Fold change cutoffs of ≥ 2.0 (RNAseq) and ≥ 1.25 (metabolomics) were employed with an FDR ≤ 0.05. n = 6 male rats per group for RNAseq and n = 7 male rats per group for metabolomics. FDR = false discovery rate.

### ZSF1-Obese HFpEF hearts displays signatures of inflammation, mitochondrial dysfunction, and downregulation of energy metabolism

Based on our physiological phenotyping results, the two-hits of obesity (i.e., metabolic syndrome) and hypertension are required for the robust development of HFpEF. Therefore, while we did examine the transcriptomic and metabolomic differences between ZSF1-Obese and WKY rats (**Supplemental Fig. 2**), we were most interested in identifying potential transcriptional and metabolic alterations revealed with the addition of obesity. A total of 5,691 genes were differentially expressed (3,123 upregulated and 2,568 downregulated; FC ≥ 2.0 and FDR ≤ 0.05) (**Fig. 3A**). Interestingly, fibrosis and inflammation were the dominant signatures based on GO enrichment analyses, including pathways related to extracellular matrix assembly, immune cell activation, phagocytosis, B cell activation and signaling, immune response, and NF-κB signaling (**Fig. 3B,C**).

**Fig. 3:**
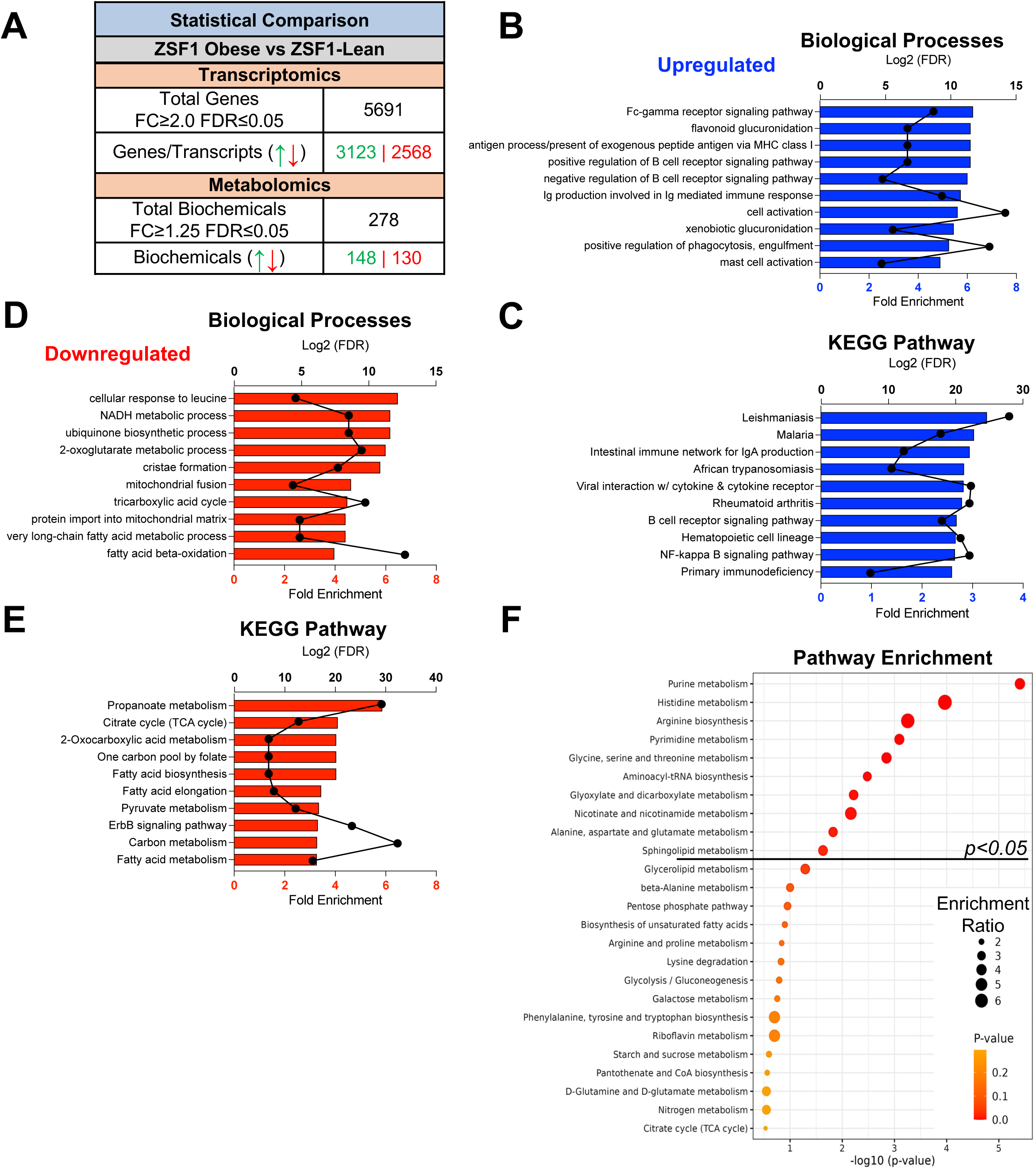
Transcriptional cardiac remodeling is dependent upon the two hits of obesity and hypertension in ZSF1-Obese HFpEF rats. RNAseq and metabolomic comparisons of hearts from ZSF1-Obese vs Lean rats. (**A**) Summary of the upregulated and downregulated transcriptional and metabolic changes. Gene ontology analysis revealing the top upregulated (**B**) biological processes and (**C**) KEGG pathways of those genes found to be differentially expressed. Gene ontology analysis revealing the top downregulated (**D**) biological processes and (**E**) KEGG pathways of those genes found to be differentially expressed. (**F**) Pathway enrichment analysis of the cardiac metabolome indicated those pathways found to be most significantly affected in ZSF1 rats due to hypertension. Fold change cutoffs of ≥ 2.0 (RNAseq) and ≥ 1.25 (metabolomics) were employed with an FDR ≤ 0.05. n = 6 male rats per group for RNAseq and n = 7 male Lean and 8 male ZSF1-Obese rats for metabolomics. FDR = false discovery rate.

GO enrichment analysis of significantly downregulated transcripts revealed suppression of key metabolic and mitochondrial biological processes (**Fig. 3D**). This included the downregulation of ubiquinone biosynthesis, cristae formation, fusion, and protein import into the matrix (**Fig. 3D**). Metabolic pathways that were downregulated in ZSF1-Obese hearts included the Krebs cycle, fatty acid metabolism, and pyruvate metabolism (**Fig. 3E**). In agreement with the transcriptomic analyses, the metabolomic signature was impacted to a greater degree than that observed with hypertension alone (i.e., Lean vs WKY; Fig. 2), with 148 metabolites that were increased and 130 metabolites that decreased in ZSF1-Obese rats as compared to Lean controls (FC ≥ 1.25 and FDR ≤ 0.05) (**Fig. 3A**). Pathway enrichment analysis revealed nucleotide and amino acid metabolism as the most impacted metabolic processes in HFpEF hearts (**Fig. 3F**); although fewer total pathways were significantly impacted, this was due to the underlying metabolic remodeling invoked by hypertension alone. In fact, several metabolites associated with pathways significantly enriched by hypertension alone (e.g., Krebs cycle) were further disrupted in the ZSF1-Obese heart, comparison of the ZSF1-Obese vs WKY in **Supplemental Fig. 2F**, which shows similarly enriched pathways in Lean vs WKY.

### Omics integration reveals transcriptional and metabolic coordination of the cardiac energetic deficit in HFpEF

To gain further mechanistic insight into HFpEF development, we next examined transcriptional changes dependent on the two-hits of metabolic syndrome + hypertension versus those independent of hypertension. Changes independent of hypertension included 795 differentially expressed genes (**Supplemental Fig 3A**), with an enrichment in processes related to the cell cycle and proliferation. This transcriptional enrichment could be associated with the meta-inflammation known to occur in HFpEF and which appears evident in the ZSF1-Obese hearts (**Fig. 3B,C** and **Supplemental Fig 3B**). Transcriptional changes dependent on both hypertension and metabolic syndrome revealed 5,544 differentially expressed genes with a significant enrichment in energy metabolism pathways and additional signatures of inflammation (**Supplemental Fig 3C**).

Merger of our omics data sets provides a more comprehensive and integrated interpretation of the remodeling occurring in HFpEF. Using a multi-omics assimilation approach, the differentially expressed transcripts and metabolites significantly altered in abundance were integrated to reveal pathways most impacted that likely contribute to disease progression. The effects of hypertension alone (Lean vs. WKY) indicated glycolysis, purine and pyrimidine metabolism, and nicotinate and nicotinamide metabolism as pathways most impacted (**Fig. 4A**). HFpEF hearts (ZSF1-Obese) had a greater impact on metabolic pathways related to ketone bodies, lipid metabolism, pyruvate metabolism, and the Krebs cycle, which was the most impacted pathway (**Fig. 4B**).

**Fig. 4:**
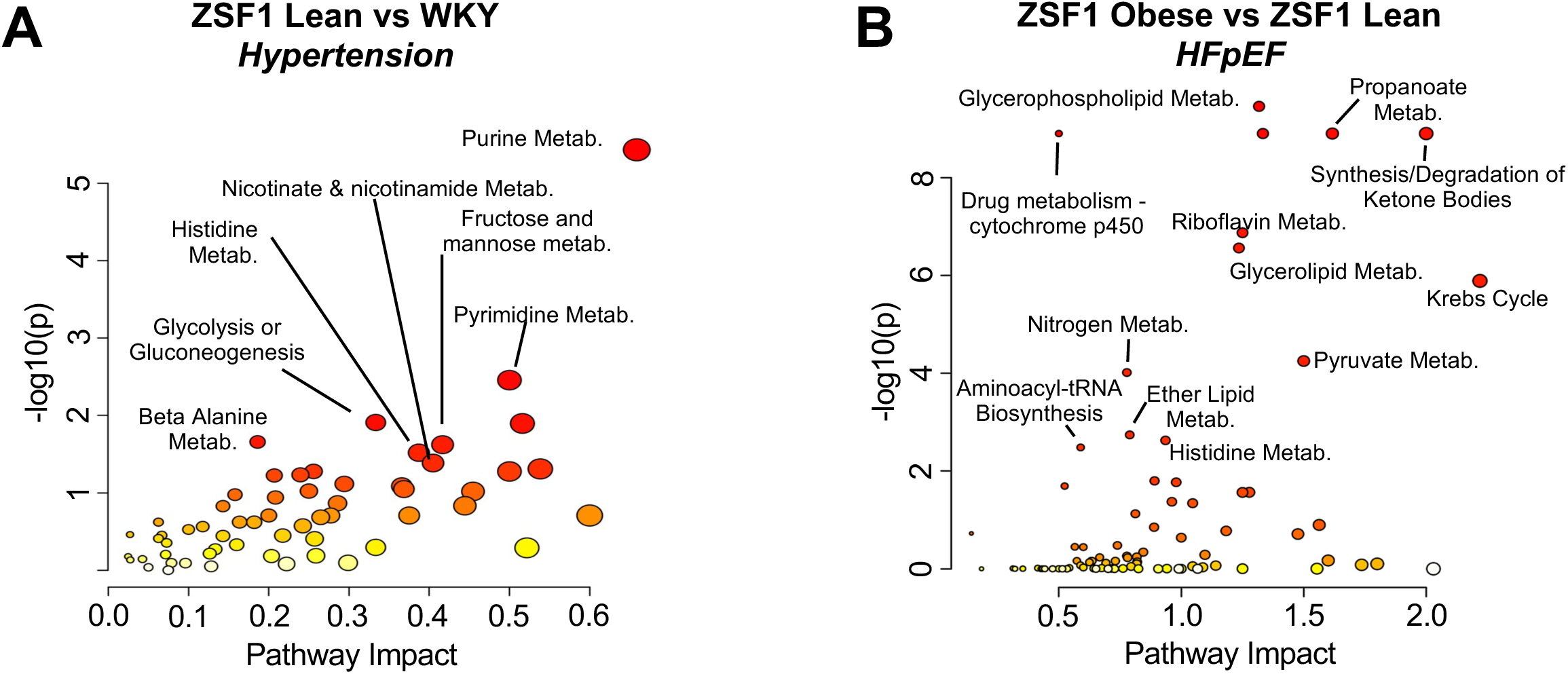
Integrated network analysis of RNAseq and metabolomic dataset reveals unique metabolic pathways impacted in HFpEF. Integrated analysis of cardiac omics datasets to identify those pathways most impacted by transcriptional and metabolic remodeling due to (**A**) hypertensive phenotype (Lean vs WKY) or (**B**) the observed HFpEF phenotype (ZSF1-Obese vs Lean). Fold change cutoffs of ≥ 2.0 (RNAseq) and ≥ 1.25 (metabolomics) were employed with an FDR ≤ 0.05. A greater pathway impact indicates a greater influence at the transcriptional and metabolic level to a given pathway. Labeled pathways had a *p-*value ≤ 0.05. n = 6 male rats per group for RNAseq and n = 7 male Lean and 8 male ZSF1-Obese rats for metabolomics. FDR = false discovery rate.

As many of the identified pathways are central to cardiac energy metabolism (i.e., glycolysis, pyruvate metabolism, Krebs cycle), we generated integrated metabolic pathway maps to better illustrate the transcriptional and metabolic changes in these pathways. The hypertensive effects (i.e., ZSF1-Lean vs. WKY) on glycolysis revealed increased expression of *Pfk* (phosphofructokinase) and *Pfkfb1* (6-phosphofructo-2-kinase:fructose-2,6-bisphosphatase), the later which generates fructose-2,6-bisphosphate, a potent allosteric activator of PFK^24^. The downstream glycolytic intermediates 3-phosphoglycerate, 2-phosphoglycerate, phosphoenolpyruvate, and pyruvate were all increased in abundance, potentially suggesting increased glycolytic activity in Lean hearts compared to WKY controls (**Fig. 5A**). This is in stark contrast to the ZSF1-Obese HFpEF heart which showed an overall downregulation of glycolytic enzymes. Obese hearts when compared to Lean had a higher PCr:ATP ratio and lower ATP:ADP ratio than Lean vs WKY, indicating a lower cardiac energy state in HFpEF. These differences were largely driven by a reduction in ATP abundance in the HFpEF heart (**Fig. 5A**). Interestingly, PCr levels were highest in the HFpEF heart, likely in part due to transcriptional downregulation of creatine kinase isoforms (i.e., *Ckm*, *Ckmt2*).

**Fig. 5:**
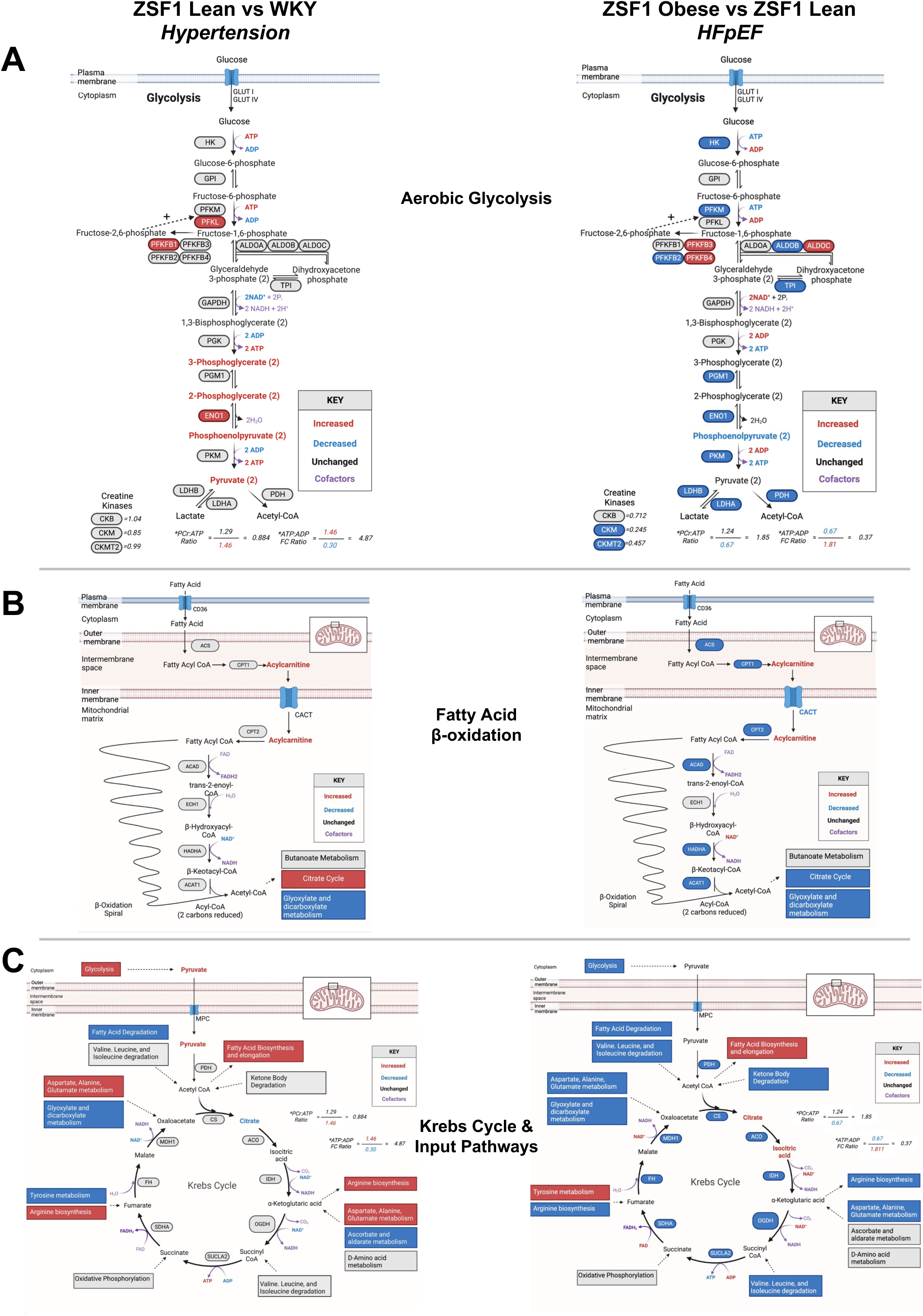
HFpEF results in the transcriptional and metabolic downregulation of pathways central to energy metabolism. Pathway maps of the transcriptional and metabolic alterations in Lean vs WKY and ZSF1-Obese vs Lean rats in energy generating pathways, specifically (**A**) aerobic glycolysis, (**B**) fatty acid oxidiation, (**C**) and the Krebs cycle. Additional pathways indicated in boxes provide an objective summary of the transcriptional and metabolic increase or decrease observed. Genes and metabolites significantly **increased** or **decreased** in expression or abundance (Fold change ≥ 2.0 (RNAseq) and ≥ 1.25 (metabolomics); FDR ≤ 0.05) are as indicated.

Hypertensive and HFpEF hearts demonstrated increased abundance of acyl-carnitines, with greater increases in the two-hit hearts (ZSF1-Obese), suggestive of decreased utilization or increased synthesis (**Fig. 5B, Supplemental Fig. 4**). ZSF1-Obese hearts also showed downregulation of key β-oxidation enzymes and transporters (e.g., *Cact, Cpt1, Cpt2, Acat1*) (**Fig. 5B**). Transcriptional repression of all Krebs cycle enzymes accompanied by increased abundance of the upstream metabolites citrate and isocitrate, suggest an overall decrease in Krebs cycle activity in Obese hearts (**Fig. 5C**).

As glycolysis and β-oxidation are central to cardiac oxidative metabolism, downregulation of their enzymes along with additional pathways capable of input to the Krebs cycle [i.e., branched chain amino acids (BCAAs), ketones, amino acids] also likely contributes to the apparent overall decrease in Krebs cycle activity and the energetic deficit of the HFpEF heart (**Fig 5 and Supplemental Fig. S5**). These integrated analyses reveal a transcriptional and metabolic signature brought upon by obesity in HFpEF, highlighting mitochondrial energy metabolism as a potential distinguishing and important feature.

### Disrupted mitochondrial ultrastructure and impaired function are evident early in HFpEF development

Due to the strong mitochondrial signature unique to HFpEF, we looked deeper and examined mitochondrial ultrastructure by transmission electron microscopy. Gross qualitative assessment of electron micrographs revealed mitochondrial cristae disorganization, with less dense cristae observed in the Lean and this progressively worsened in Obese hearts **(Fig 6A)**. Quantitative mitochondrial morphological analyses indicated no difference in mitochondrial number per cardiomyocyte area but a decrease in the total mitochondrial area, indicating smaller mitochondria in Obese hearts (**Fig. 6B**). Damaged or fragmented mitochondria typically assume a smaller and more rounded morphology^32,33^, this was evident by a reduction in Feret’s diameter and an increase in the circularity index in ZSF1-Obese HFpEF cardiomyocyte mitochondria (**Fig. 6C**). Strikingly, obese hearts displayed a significant increase in lipid droplets (LDs), which localized adjacent to interfibrillar mitochondria (**Fig. 6A**). Quantification of LDs revealed a significant increase in number and size exclusively in Obese hearts (**Fig. 6D,E**). Because LDs strongly associated with interfibrillar mitochondria, we quantified mitochondria-LD interactions which indicated an increase in the total number of mitochondria-LD contacts as well as the length of mitochondrial and LD membranes in close association with one another (**Fig. 6F**). Lastly, sarcomeric length was increased in ZSF1-Obese cardiomyocytes, likely a consequence of increased preload (i.e., diastolic dysfunction) and LV dilation observed in the HFpEF heart (**Fig. 6G**).

**Fig. 6:**
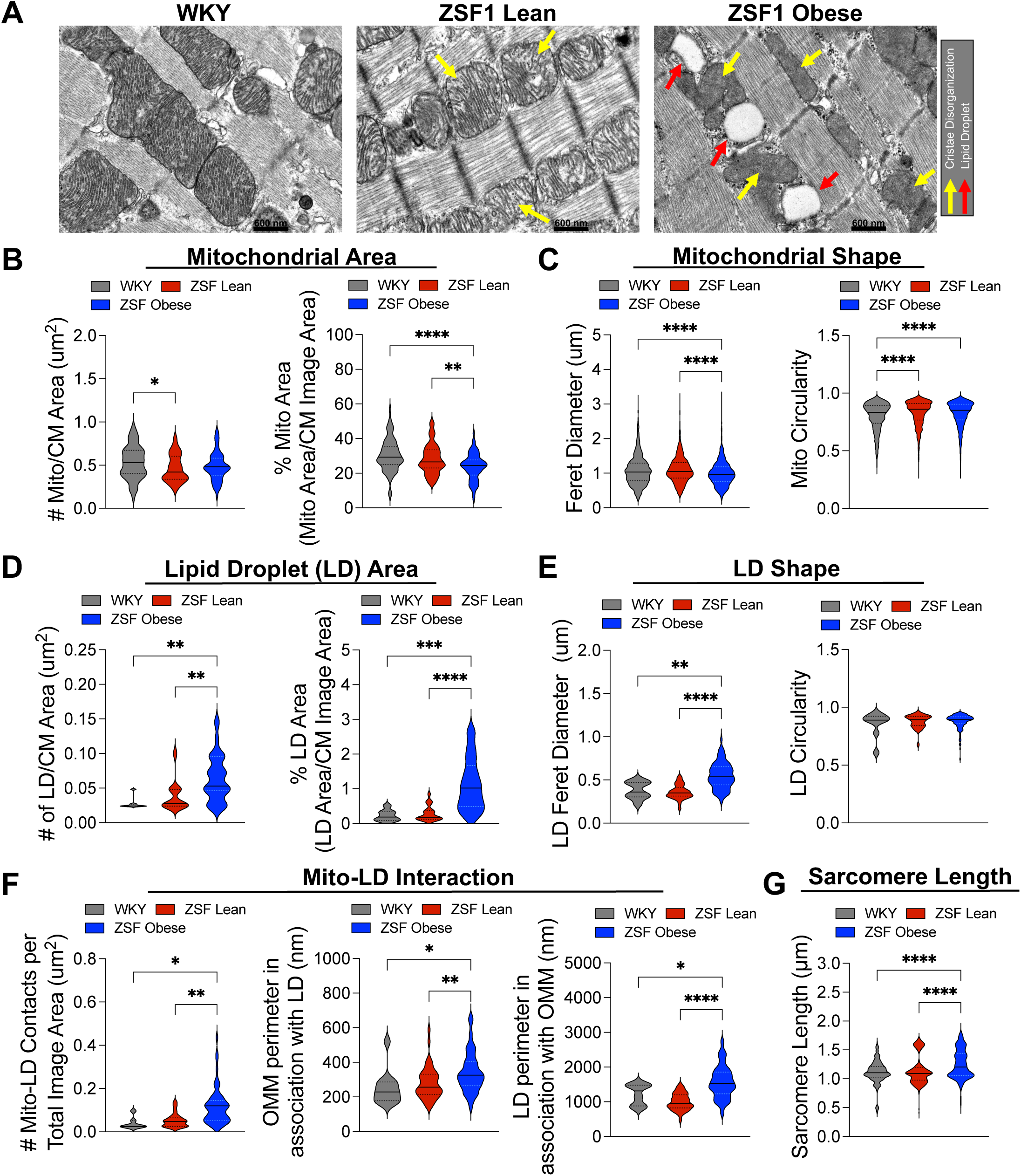
Mitochondrial ultrastructural remodeling in the HFpEF heart is characterized by a decrease in mitochondrial content, cristae disorganization, and lipid droplet association. Transmission electron micrographs of cardiomyocyte mitochondrial and lipid droplet (LD) ultrastructure. (**A**) Representative images of WKY, Lean, and ZSF1-Obese cardiomyocyte ultrastructure indicating cristae disorganization (**yellow arrows**) and lipid droplet accumulation and interaction with mitochondria (**red arrows**). (**B**) mitochondrial area (i.e., content) quantified by the number of mitochondria per image area and the percent area of mitochondria to total area. (**C**) Mitochondrial shape quantified by the Feret’s diameter and circularity index. (**D**) Quantification of LD area (i.e., content) quantified by the number of LDs per image area and the percent area of LDs to total area. (**E**) LD shape quantified by the Feret’s diameter and circularity index. (**F**) Analysis of mitochondrial-LD interaction quantified by the number of mito-LD contacts per image area, the outer mitochondrial membrane (OMM) perimeter in contact with an LD, and the LD perimeter in association with the OMM. (**G**) Determination of sarcomeric length measured from z-line to z-line. n = 4 male rats per group, mean ± SEM. One-way ANOVA with Holm-Sidak’s post-hoc test, *p ≤ 0.05, **p ≤ 0.01, ***p ≤ 0.001, ****p ≤ 0.0001.

With notable mitochondrial ultrastructural changes, we next examined mitochondrial function via respiratory activity and calcium handling assays. Determination of citrate synthase activity, a gold standard for assessing mitochondrial abundance, was decreased in both Lean and Obese rats (**Fig. 7A**). Both Lean and Obese cardiac mitochondria displayed lower overall respiratory rates as compared with WKY controls (**Fig. 7B,D,F**). In the presence of pyruvate + malate (complex I) or succinate + rotenone (complex II), mitochondria from ZSF1-Lean and –Obese hearts showed a significant reduction in state 3 respiration (**Fig. 7C,E**); fatty acid supported state 3 respiration (palmitoyl-l-carnitine) also trended lower, but did not reach statistical significance (**Fig. 7G**). Complex I respiratory control ratio (RCR) was reduced in the Lean hearts, indicating reduced coupling of oxygen consumption to ATP production, and surprisingly this was improved in the Obese hearts when compared to the reduction in Lean (**Fig. 7C**). No differences were observed for complex II RCR, FAO RCR, or State 4 rates (**Fig. 7E,G and Supplemental Fig. 6A**).

**Fig. 7:**
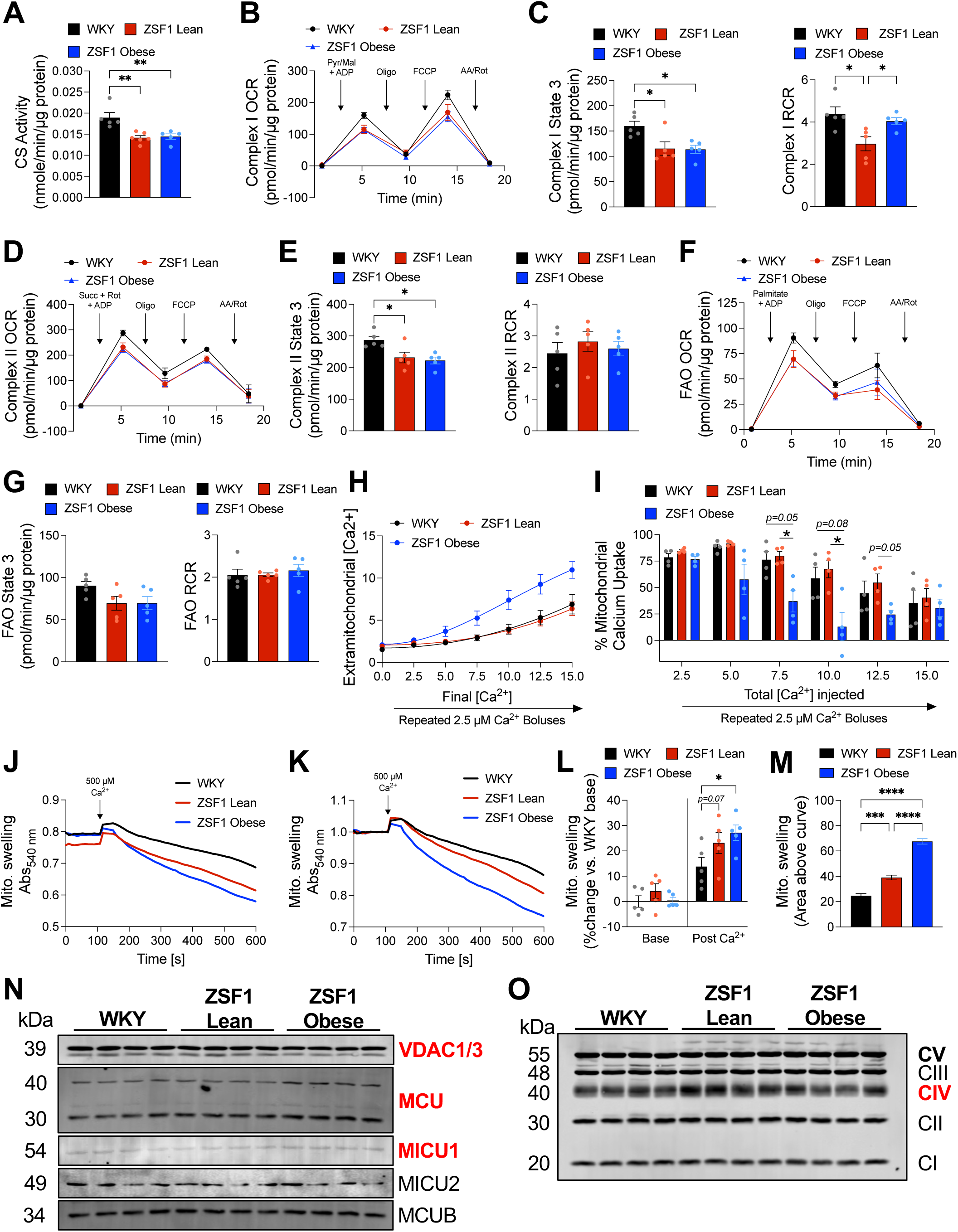
Mitochondrial dysfunction characterized by impaired respiratory activity and disrupted calcium handling is a key feature of the HFpEF heart. Functional assessment of the mitochondrial function in the HFpEF heart. (**A**) Determination of mitochondrial content assessed by citrate synthase activity. Interrogation of mitochondrial respiratory function by assessing oxygen consumption rates (OCR) of (**B,C**) complex I (pyruvate + malate)-specific substrates, (**D,E**) complex II (succinate)-specific substrate + rotenone (complex I inhibitor), and (**F,G**) fatty acid oxidation capacity (palmitoyl-L-carnitine): state 3 (substrate-mediated) oxygen consumption and the respiratory control ratio (RCR) providing an index of oxygen consumption to ATP-production coupling. (**H,I**) Mitochondrial calcium uptake in response to repeated 2.5 µM boluses. Mitochondrial swelling in response to a 500 µM bolus, displayed as both (**J**) uncorrected and (**K**) normalized prior to calcium addition. Quantification of mitochondrial swelling indicated by (**L**) percent change to WKY baseline and (**M**) area above the curve. Immunoblotting of (**N**) VDAC1/3 and components of the mitochondrial calcium uniporter and (**O**) subunits of ETC complexes. Proteins differentially expressed in protein abundance are indicated in **red**. n = 5 male rats per group (A-G), n = 4 male rats per group (I-O), mean ± SEM. One-way ANOVA with Holm-Sidak’s post-hoc test, *p ≤ 0.05, ***p ≤ 0.001, ****p ≤ 0.0001.

Mitochondrial calcium uptake, is intricately linked to bioenergetics^34^ and at high levels induces mitochondrial dysfunction. Indeed, HF is associated with mitochondrial calcium overload^26,27,34,35^. We isolated mitochondria from ZSF1-Obese hearts and subjected them to repeated 2.5 µM Ca^2+^ boluses. Interestingly, ZSF1-Obese cardiac mitochondria failed to uptake Ca^2+^ as demonstrated by the accumulation of Ca2+ in the bath (i.e., extramitochondrial) (**Fig. 7H,I** and **Supplemental Fig. 6B**). This is suggestive of mitochondria that are either already calcium-overloaded or that have downregulated mitochondrial calcium uniporter activity. Mitochondrial swelling, an indicator of susceptibility to mitochondrial permeability transition, was increased in Lean hearts while swelling of ZSF1-Obese HFpEF mitochondria occurred faster and to a greater extent than both WKY and Lean mitochondria (**Fig. 7J-M and Supplemental Fig. 6C**). Protein expression via immunoblotting of proteins involved in mitochondrial Ca^2+^ handling revealed a significant increase in both the 30 kDa and 40 kDa MCU isoforms and in the MCU gatekeeper, MICU1, exclusively in ZSF1-Obese HFpEF cardiac mitochondria when normalized to a mitochondrial loading control, ATP synthase (i.e., complex V) (**Fig. 7N,O and Supplemental Fig. 7**). VDAC1, which is also involved in mitochondrial calcium homeostasis^34^, was significantly decreased in ZSF1-Obese mitochondria (**Fig. 7N**). Collectively, our results indicate significant remodeling of the mitochondrial ultrastructure, accumulation of cardiomyocyte lipid droplets, dysfunctional respiratory capacity, and dysregulated calcium handling all of which underlie and likely contribute to the gross metabolic dysregulation and subsequent cardiac dysfunction observed in the HFpEF heart.

## DISCUSSION

In the present study, we characterized and identified the underlying molecular changes associated with the HFpEF phenotype in a robust preclinical rat model. Key findings include:

- Metabolic syndrome/obesity is a principal driver of HFpEF
- Transcriptional and metabolic remodeling in HFpEF is characterized by the upregulation of inflammation and downregulation of energy metabolism
- Mitochondrial ultrastructural and functional remodeling underlie the HFpEF phenotype and likely is an early contributor to cardiac dysfunction.
- The HFpEF heart displays significant intramyocardial lipid accumulation (huge increase in lipid droplet size, number, and association with mitochondria).

### Impact of the additive hit of metabolic syndrome and obesity

The ZSF1-Obese rat model recapitulates the multifactorial clinical features that distinguish HFpEF. Importantly, our study validates the notion that “two-hits”, hypertension and metabolic syndrome/obesity are necessary for the development of HFpEF, similar to the L-name + HFD murine model^11^ and in agreement with human HFpEF populations which are typically obese with vascular dyfunction^36,37^. A strength of our study is the inclusion of the WKY non-hypertensive control as this allowed us to examine transcription, metabolic, and functional changes that are dependent and independent of these “two-hits”. While hypertension alone resulted in cardiac metabolic remodeling, the addition of metabolic syndrome resulted in more drastic metabolic remodeling which was associated with an energetic deficit and mitochondrial abnormalities, both in ultrastructure and function. Similarly, only 152 transcription changes were exclusively dependent upon hypertension, whereas 795 transcripts independent of hypertension and 5,544 transcript changes were dependent on both metabolic syndrome and hypertension. While other models of HFpEF have been proposed, namely models of Western diet feeding^9^, angiotensin II/phenylephrine (ANGII/PE) infusion^8^, and senescence-accelerated aging (i.e., SAMP/SAMPR mice)^10,12^, and while pathology associated with these models may be multifactorial, these models do not contain two independent hits. Furthermore, long-term Western diet feeding progresses to HFrEF, which rarely occurs in humans^38,39^, and ANGII/PE and SAMP/SAMPR models lack metabolic syndrome and/or hypertension. While these models are likely suitable for the study of diastolic dysfunction, this is distinct from the multifaceted, multi-organ nature of HFpEF. Thus, the ZSF1-obese rat model serves as an excellent preclinical model to study cardiometabolic HFpEF.

Identifying therapeutic targets for HFpEF has proven difficult due to the combinatorial etiologies that contribute to the syndrome. Recently, SGLT2 inhibitors (SGLT2i), which act to block glucose reabsorption in the kidney, have proven efficacious and safe in reducing cardiovascular events in animal models^40,41^ and in HFpEF trials^42,43^. SGLT2i have a minimal impact on hypertension yet result in profound weight loss and normalization towards glucose homeostasis^40,44,45^. These results are directly in line with our findings and overall conclusion that the primary driver of HFpEF is the metabolic syndrome component, which is further exacerbated in the Obese-vs Lean-ZSF1 rats. Adjunctive therapy of SGLT2i + a hydrogen sulfide donor (H_2_S, a well-studied cardioprotective agent) in our ZSF1-Obese HFpEF model was shown to be efficaciously superior to either treatment alone^19^. This is interesting as H2S has been shown to modulate metabolism^23^ and preserve mitochondrial integrity^46^. While we observed numerous transcriptional changes independent of hypertension, most of the transcriptional remodeling was dependent on both hits, thus, treatments aimed at targeting metabolic syndrome alone will likely be insufficient for long-term efficacy.

Our study provides a roadmap for the discovery of novel mechanisms driving HFpEF progression and provides a data set which can be correlated to the remodeling observed in human HFpEF^47^. Targeting of HDAC6 in a HFpEF mouse model was recently shown to be as efficacious as SGLT2i^41^. This is of note as the mechanisms of SGLT2i cardioprotection remains unclear as mice with global loss of SGLT2 are protected from HF with SGLT2 blockade^48,49^, indicating SGLT2 inhibitors likely have an off-target mechanism of action.

### Remodeling of Energy Metabolism

An important observation from our metabolomics dataset is a drastic change in the energy state of the ZSF1-Obese heart. HFpEF hearts displayed a greater PCr:ATP ratio (1.85 vs 0.884) and lower ATP:ADP ratio (0.37 vs 4.87) as compared to hypertension alone. This was driven by a significant reduction in ATP in the HFpEF heart. This may, in part, be associated with an inability to liberate PCr stores, as all creatine kinase isoforms were reduced in ZSF1-Obese rats. This indicates that the HFpEF heart is energy starved, as compared to the ZSF1-Lean control (hypertension alone). The ZSF1-Obese heart also displayed gross metabolic remodeling of pathways associated with energy metabolism, discussed hereafter.

#### Glycolysis

We observed an increase in the expression of *Pfkl* and of *Pfkfb1*, activators of aerobic glycolysis, and downstream metabolites of glycolysis (i.e., 3-PG, 2-PG, PEP, and pyruvate) were also found to be increased, suggesting increased glycolysis in the hypertensive heart. In contrast, nearly all glycolytic enzymes were decreased in abundance in the Obese HFpEF heart. These results are in agreement with results from the Kass Lab which also found a reduction in protein expression of these same glycolytic enzymes in human HFpEF endomyocardial biopsies; however, they only detected decreased abundance of the upstream glycolytic metabolites G6P and F-1,6-BP while we observed a decrease in the downstream metabolite PEP^50^. Glucose oxidation is also likely decreased as we and others have shown changes in the abundance of pyruvate, PDH, MPC1, and PDK4^50,51^, with direct measurements of reduced glucose oxidation performed in the working heart^52^. Both in the ZSF1-Obese and human HFpEF heart, changes in the pentose phosphate pathway (i.e., purine and pyrimidine metabolism) were identified, indicating disruptions to ancillary biosynthetic pathways; these ancillary pathways are known to contribute to cardiac remodeling^53,54^, highlighting that their role in HFpEF is an area worthy of investigation. Collectively, these studies suggest a significant downregulation of glycolytic metabolism and changes in ancillary pathways in the HFpEF heart.

#### Fatty Acids

Fatty acids contribute the largest percentage to cardiac energy production, thus loss of oxidative capacity in the failing heart would be detrimental to energy metabolism and cardiac function. Our results indicate a significant impairment in fatty acid oxidation (FAO), with associated FA metabolic processes among the most downregulated in the HFpEF heart, including key enzymes in transport and processing (e.g., *Acs*, *Cpt1*, *Cact*, *Acad*, *Acat1*). Dysregulation of genes involved in fatty acid and oxidative metabolism seems a conserved signature, as similar findings to ours have been shown in other murine models^41,55^ and human HFpEF populations^51^; however, some studies have reported an increase in FAO transcripts^56^. While there is a discrepancy in gene expression among different studies, a proteomic study of HFpEF samples, from the same group that reported an increase in FAO and OXPHOS transcripts, found an overall decrease in protein abundance^57^, a reminder that transcript and protein abundance often do not correlate in pathology.

In agreement with a decrease in FAO, our results indicate reduced abundance of short chain acyl-carnitines and increased medium and long-chain acylcarnitines, potentially suggesting inefficient oxidation. Medium and long-chain acylcarnitines were decreased in human HFpEF and the expression of FAO genes were also decreased^51^; whether this discrepancy in findings is simply due to sampling of the right ventricle versus the left ventricle remains to be determined. Nonetheless, Krebs cycle intermediates are lower in human HFpEF^51^ and we demonstrate reduced utilization of fatty acids by mitochondria isolated from HFpEF hearts. While these collective results suggest impairments in fatty acid utilization, palmitate oxidation measured in the isolated working heart was increased in the mouse L-name + HFD mouse model^52^; thus, more work is needed to define how HFpEF remodels cardiac fatty acid metabolism.

#### Ketones, BCAAs, and Amino Acids

The potential for alternative fuel sources to contribute to cardiac energetics has become more appreciated. Our integrated network analysis approach identified the synthesis and degradation of ketone bodies as highly impacted in HFpEF. We observed an increase in 3-hydroxybutyrate and a corresponding decrease in key ketone catabolic enzymes (i.e., *Bdh1*, *Oxtc1*, *Acat1*), suggestive of reduced utilization. In a murine model of HFpEF, BDH1 protein abundance is reduced with a corresponding trend of decreased oxidation rates^52^. In HFrEF, myocardial uptake, oxidation, and expression of BDH1 increases 2-to 3-fold^58–60^, which is greater than predicted rates in HFpEF^59^. This divergence in ketone body oxidation between HFrEF and HFpEF may provide insight into differential substrate/fuel treatment strategies. For example, while increasing circulating ketones through the diet appears to provide beneficial effects in HFrEF^61^, whether this would be beneficial in HFpEF has not been explored. Another potential target could be HMGCS, which we found upregulated in the ZSF1-Obese HFpEF heart and is generally known to be involved in ketone synthesis; thus, whether impaired ketone oxidation is due to competing synthesis mediated by HMGCS presents an interesting inquiry.

The branched chain amino acids (BCAAs) leucine, valine, and isoleucine have been proposed as an alternative fuel source for the heart and suppression of BCAA oxidation has been implicated in heart failure^62,63^. While we observed a global suppression of BCAA oxidation genes, the downregulation of the nodal BCAA catabolic enzyme, branched-chain α-keto acid dehydrogenase complex (*Bckdh*), agrees with data in human HFpEF^51^. Previous reports suggest BCAAs accumulate in the human HFpEF heart, suggesting decreased oxidation^64^. However, contributions of BCAAs to energy production are likely minimal^59,62,65,66^ and the activation of cardiac BCAA oxidation does not provide energetic or functional benefit in models of HFrEF^67^. Thus, while BCAA oxidation seems downregulated, targeting this pathway in HFpEF may not prove effective.

Our integrated pathway maps identified the downregulation of numerous other amino acid pathways at the level of transcription and/or metabolite abundance. The observed changes in global amino acid metabolism could be related to the increased proteolysis that occurs in the failing heart^59^. Many of these amino acids and represented pathways have yet to be explored, providing experimental opportunities to generate new hypotheses. For example, we observed a reduction in arginine metabolism, which when given as an oral supplement to HFrEF patients proved beneficial^68^; whether similar benefits could be obtained in HFpEF patients is worth exploring.

### Impact on Mitochondrial and Lipid Droplet Structure and Function

*Ultrastructural changes* – Mitochondrial dysfunction is a hallmark of HF^69–71^ and our transcriptomic and metabolic signatures implicates the derangement of several mitochondrial processes in the ZSF1-Obese heart. Downregulation of biological processes related to cristae formation and mitochondrial fusion were confirmed by ultrastructural remodeling characterized by the disruption and near disappearance of cristae and overall smaller and more rounded mitochondria, indicating that the fission-fusion balance is perturbed. Alterations in mitochondrial shape and cristae density can greatly impact the localization, structure and function of the OXPHOS system, impairing cellular and mitochondrial metabolism^32,33,72^. While we observed no change in total mitochondrial number, mitochondrial area was significantly reduced in HFpEF, likely because the smaller size of individual mitochondrions. Interesting, mice treated with the SGLT2i empagliflozin demonstrated an increase in mitochondrial area per cardiomyocyte area^40^. Recently, TEM of human HFpEF cardiomyocytes revealed no change in mitochondrial area but significant cristae derangement which was most observable in patients presenting with obesity^57^.

There exists a high correlation between myocardial adiposity and diastolic dysfunction^73,74^. Cardiac MRI of HFpEF, HFrEF, and non-failing patients revealed significant intramyocardial fat only in HFpEF^75^. We observed the accumulation of LDs in HFpEF hearts which was also recently seen in HFpEF patients via TEM imaging^57^. LDs act as an energy storage depot and are involved in transferring stored FAs to mitochondria for energy production. However, transcriptional downregulation of fatty acid oxidation machinery and the structural remodeling likely limit utilization, thus promoting storage and LD accumulation. While we observed greater mito-LD interactions in HFpEF, the interpretation of this result is confounded by the fact that few, if any LDs were observed in control hearts. To overcome this limitation, we examined the expression of known proteins that act as tethers to support mitochondria and lipid droplets approximation. Perilipin 5 (PLIN5), a LD protein reported to tether them to mitochondria^76,77^, was downregulated 3-fold in our HFpEF hearts. Loss of PLIN5 decreases mito-LD interactions and oxidative metabolism, whereas overexpression increases these interactions^78^. Similarly, we noted a downregulation of *Miga2*, another mito-LD tether involved in lipid metabolism and mitochondrial fusion^79,80^. Whether disruption of these tethers could play a role in HFpEF is unknown, but it’s striking that these LD proteins were downregulated in the context of massive LD biogenesis.

*Mitochondrial Dysfunction* – Mitochondrial respiratory capacity was significantly impaired in both the pre-HFpEF (hypertensive) and HFpEF (hypertension + metabolic syndrome) heart, suggesting that while mitochondrial dysfunction is a key feature of HF, it is not necessarily unique to HFpEF. What is unique in the HFpEF heart is impaired mitochondrial calcium handling. Mitochondrial protein expression of MCU and MICU1, components of the mitochondrial calcium uniporter, were increased exclusively in the HFpEF heart. This could be a compensatory change to increase calcium-dependent activation of mitochondrial dehydrogensases to increase Krebs cycle flux and mitochondrial energetics. However, as previously reported by our group and others, while initially compensatory these expression changes in uniporter components turns maladaptive with chronic stress (refs). In a mouse model of HFpEF, SGLT2i treatment improved HFpEF-mediated Ca^2+^ reuptake by the sarcoplasmic reticulum and rescued mitochondrial respiratory function^40^; however, whether improved reuptake was a consequence of improving mitochondrial Ca^2+^ buffering, improved energetics, and/or enhancing SERCA activity was not tested. Similarly, treating HF with a pan HDAC inhibitor (SAHA) decreased acetylation of proteins involved in oxidative metabolism, improving mitochondrial oxidative phosphorylation^81^. Whether HDAC inhibition plays a similarly protective role in HFpEF remains to be investigated.

### Study Limitations

There are several limitations to our study. Our results do not test a specific mechanism or hypothesis but provide an integrated systems biology approach to allow for the discovery of potentially important pathways and mechanisms contributing to HFpEF. The current study also exclusively utilized male rats, partly due to the high-cost of acquiring a sufficient number of female rats to maintain power in our dual-control study. However, a recent study that exclusively utilized ZSF1 female rats reported similar findings related to the mitochondria^40^, suggesting conserved mechanisms of action between sexes. Adjusting for sex in a human HFpEF RNAseq study, importantly, did not affect pathway enrichment^56^. Nonetheless, as the prevalence of HFpEF is slightly greater in females than males, we understand and acknowledge the importance of potential for sex differences in molecular pathways. Current work in our labs is exploring whether similar functional, metabolic, transcriptional, and mitochondrial remodeling is found in female HFpEF or whether sex distinguishes between remodeling pathways and targets. Also, aging is a critical risk factor for HFpEF and is not accounted for in our study. As we have shown in a large animal model of diastolic dysfunction, aging alone significantly alters the cardiac transcriptome and metabolome^7^, making it difficult to tease out pathway changes due to disease progression, aging, or both. Lastly, it is evident that metabolic syndrome and obesity are primary drivers of HFpEF development, thus understanding the systemic changes at peripheral tissues is critical to complete our understanding of HFpEF pathophysiology. In a large-animal model that recapitulates several clinical features of HFpEF and diastolic dysfunction, we found skeletal muscle to have distinct transcriptional and metabolic signatures that were accompanied by mitochondrial dysfunction^7^, and similar findings have been noted in human HFpEF skeletal muscle^82^. Similar approaches have also been performed to identify potential candidates for interorgan crosstalk between the liver and heart in HFpEF^83^. Investigating peripheral tissues and potential interorgan communication likely will yield novel and meaningful insights to understand HFpEF development and progression.

## CONCLUSIONS

In summary, the results presented here demonstrate the power of applying integrated omics technologies to lead to the design of functional experiments to test specific hypotheses and discover novel therapeutic targets. The ZSF1-Obese rat model recapitulates the clinical characteristics of human HFpEF and shares many of the same transcriptional, metabolic, and mitochondrial remodeling as seen in patients. Our findings provide a wealth of data that are likely to reveal novel metabolic pathways and molecular targets which will hopefully allow for the discovery of new therapeutics to treat HFpEF.

## Acknowledgements

AAG, DJL, and JWE were involved in conception and design. AAG, KL, RBG, YT, CCF, JD, ZL, HX, MPL, NL, MC, TS, TTG, DJL, and JWE were involved in data collection, analysis, and interpretation. AAG, DJL, and JWE were involved in drafting the manuscript and revision of the manuscript. All authors were involved in final approval of the manuscript submitted. We would also like to thank Shannon Modla and Jean Ross in the University of Delaware Bio-Imaging Center for assistance in tissue processing for TEM and Gyorgy Csordas and Timothy Schneider at Thomas Jefferson University Mito Care Center for assisting in TEM image acquisition.

